# Poxvirus targeted by RFdiffusion peptide-binders

**DOI:** 10.1101/2025.05.14.654163

**Authors:** J. Coll

**Affiliations:** Department of Biotechnology. Centro Nacional INIA-CSIC. Madrid, Spain

**Author notes:** Corresponding author & (JC).

**Keywords:** F13L, vaccinia, VACV, monkeypox, MPXV, RFdiffusion, protein trained networks, peptide-binders, cyclized

## Abstract

Peptide-binders were generated by computational RFdiffusion to the F13L-homodimer interface sequences of poxviruses such as vaccinia and monkeypox. Compared to small-drug screenings and/or co-evolutions, peptide-binders have the advantages of higher affinities and easier chemical / recombinant synthesis. Among the hundreds of 20-30-mer peptide-binders randomly generated by RFdiffusion, some targeted vaccinia (VACV) F13L homodimer interface predicting low nanoMolar affinities. To improve its physiological stability, additional peptide sequences were computationally generated by cyclization/ hallucination. The resulting *de novo* cyclic peptide sequences predicted picoMolar affinities not only targeting the VACV F13L homodimer interfaces but also inner cavities previously identified by small-drug dockings to Tecovirimat-resistant monkeypox (MPXV) mutants. Because their targeting to numerous highly-conserved amino acids among poxviruses, and improved physiological stability against proteases, cyclic peptide-binders may be more adequate against resistant mutants of VACV and MPXV poxviruses.

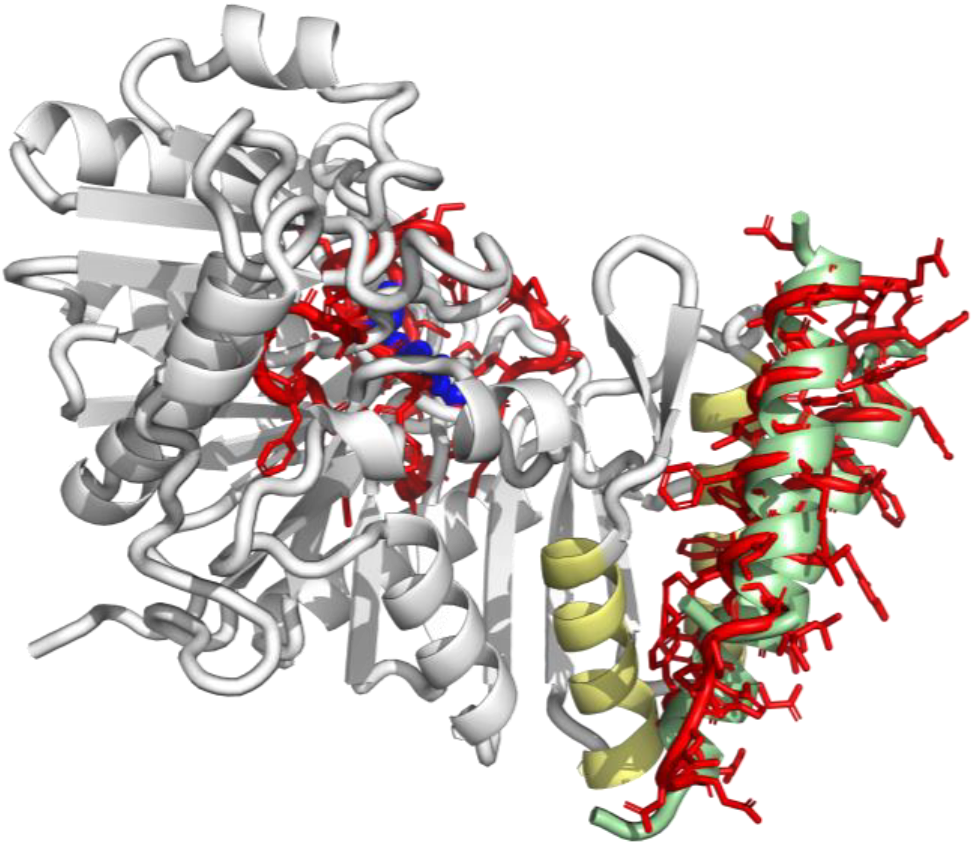

## The F13L protein of poxviruses

F13L is the most abundant membrane protein of poxviruses^1, 2^. F13L is a 372 amino acid dipalmitoylated (^185^CC^16^) protein that co-localizes with transmembrane-anchored B5R/B6R ^3-5^ poxviral glycoproteins^6^. Their deletions^7^ cause inhibition of wrapping, and reduction/attenuation of extracellularly released poxviral particles^8, 16-19^. F13L possesses main phospholipase-like (^312^NxKxxxxD)^9, 10^, and mutation-sensitive (^253^YW) ^11^ motifs.

Approved by the FDA, Tecovirimat (ST-246) is an small molecule of reference that inhibits poxvirus replication by binding F13L with a high affinity of ∼ 20 nM^20-23, 29^. Tecovirimat blocks membrane wrapping of most poxviruses including vaccinia (VACV), monkeypox (MPXV), variola, smallpox, cowpox, and camelpox, among others^12^. Tecovirimat reduces immunoprecipitation of F13L-B5R/B6R complexes^11^ and changes their intracellular membrane co-localizations^13^. Despite its demonstrated poxvirus inhibitory activities, F13L-Tecovirimat complexes have only been recently solved by crystallography (9fhs.pdb released in march of 2025). The VACV F13L shows an homodimer structure^14^ where Tecovirimat best fits the interface between chains A and B surrounded by 4 α-helices (2 α-helices per monomer)(**Figure 2A, red spheres**). These homodimer model and Tecovirimat location may explain the previous discordances between the Tecovirimat binding and its lower docking affinities.

Nevertheless, there are not yet any 3D structures solved for possible F13L-B5R/B6R complexes.

**Table 1.**
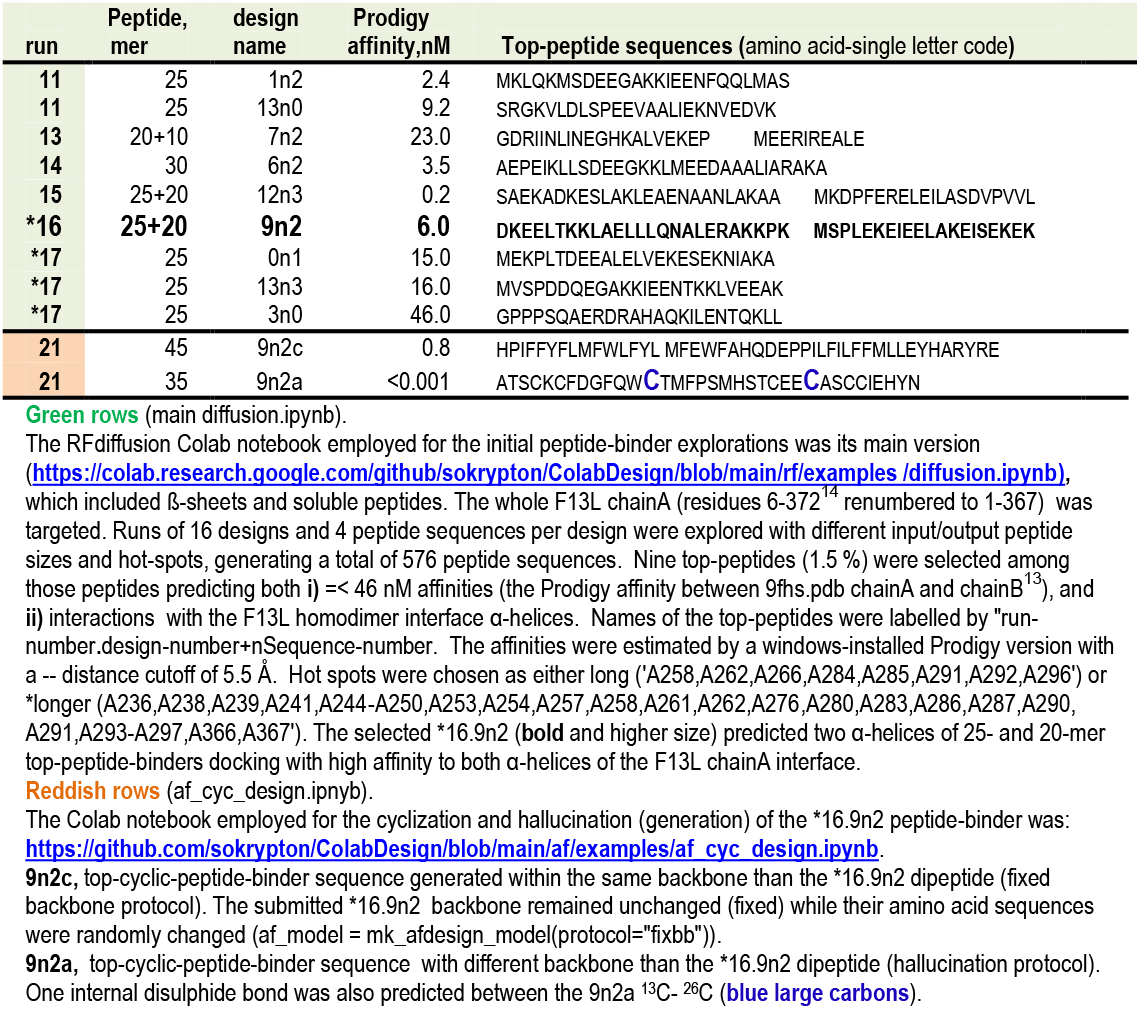
Summary of RFdiffusion runs: criteria and top-peptide results.

Before solving the homodimer structure, different authors based on their F13L alphafold monomeric VACV and/or MPXV models, suggested that Tecovirimat docking could conformationally disturb the phospholipase motif to inhibit its activity through docking/binding to nearby inner cavities ^15-18^.

Our own previously reported docking attempts targeting the MPXV F13L monomer alphafold models of Tecovirimat-resistant mutants by ***D****ata****W****arrior* ***B****uild* ***E****volutionary* ***L****ibrary* (DWBEL) successfully predicting affinity improvements on other protein/ligand pairs ^19,20^,21,22,23,24,25,26^,27,28,29,^confirmed such monomeric inner cavity hypothesis. Although large amounts of drug-like compounds could be generated by co-evolutionary docking to the inner cavities of MPXV resistant mutants with improved affinities (docking residues mapping to 52-284 + 312-338 residues)^30^, the low docking affinities of Tecovirimat to those sites were unexplained. Furthermore, despite filtering hundreds of known toxicities^2-5^, their often complex chemical synthesis and possible additional physiological side-effects remained.

Most abundant Tecovirimat-resistant poxvirus mutants have been mapped to highly conserved regions among poxviruses^3, 4^ located at the carboxy-terminal F13L domains (240-310 residues) ^28, 29, 32,30^. The isolated Tecovirimat-resistant mutants were most abundantly located before the canonical phospholipase-like motif and included the two α-helices at the F13L homodimer interface (255-268+284-299 residues)^13^.

New F13L docking backbones are needed, since no other potent inhibitors targeting Tecovirimat-resistant poxviruses such as those detected on MPXV viruses have been yet reported, despite numerous research efforts^31^.

Among other possibilities, the application of computational diffusion models to generate possible peptides inhibiting F13L activity is a recent opportunity to explore. Computational diffusion models previously developed for image generation, have been recently applied to neural networks trained by deep-learning and transformers on how to construct novel protein/peptide structures^10,32-40^. Diffusion algorithms initially diffuse protein structures with random noise and reverse/denoise them with the help of trained networks to generate *de novo* proteins/peptides. For instance, **R**osetta **F**old (RFdiffusion) have been previously employed to generate *de novo* protein-binders of ∼100 amino acids that successfully targeted snake venom 3Ftx loop III of α-neurotoxins and cytotoxins^41^. Such *de novo* peptides *in vitro* specifically neutralized their corresponding venoms and protected mice from lethal challenges. Most recently, smaller peptide-binders of ∼40 amino acids further increasing their affinities have been also generated by fine-tuning RFdiffusion of consensus snake venom sequences by introducing the new algorithms including ß-sheet, soluble-peptide and dipeptide contigs^42^.

Similar RFdiffusion algorithms have been explored here to search for more suitable alternatives to Tecovirimat, favouring their synthesis and increasing their affinities. For that, the recently solved F13L chainA homodimer interface of the VACV/MPXV amino acid sequence homodimer^13^ were targeted. The objective was to generate and select new peptide-binders to chainA that could compete with the binding affinites to chainB to disrupt the homodimer structure and inhibit F13L activity. Because of their high probabilities to interfere with the homodimer F13L interface and/or inner docking cavity, some of the cyclic/hallucinated *de novo* peptide-binder sequences generated here are proposed here for experimental work (**SupplementaryMaterials / GraphycalAbstract.pdb**).

The final described predictions may be limited in their experimental applications because of the particular F13L chainA model targeted to derive the cyclic/hallucinated top-peptide-binders, and/or the particular RFdiffusion criteria selected for these explorations. Therefore, the top-peptide-binder candidates will require additional experimental validation. Despite their possible practical limitations, these new peptide-binder strategies may provide some examples for additional explorations into the anti-poxvirus F13L peptide space.

## Computational results

### Peptide-binders to the F13L chainA

The F13L amino acids participating on chainA / chainB interaction interfaces of the VACV homodimer^14^ were studied first. The amino acid residues of F13L chainA contacts to =< 5.5 Å of chainB were identified by Prodigy and then explored in PyMol. To use for RFdiffusion random runs, the identified amino acid contacts were classified in short, long and longer hot-spots (containing 5, 8 and 34 amino acid residues, respectively).

Since preliminary random run results discarded the use of short range hot-spots by their low affinities or wrong targetings, the long and longer ranges were further explored by 6 RFdiffusion independent runs using preferences for ß-sheet and soluble peptides (**Table 1**). To facilitate the analysis of the abundant files generated from each RFdiffusion random run while increasing their probabilities to find new peptide-binder backbone designs and sequences, each of the run input/output were optimized to 16 designs, 4 sequences per design.

Because poly-H tails were added to the crystallized recombinant F13L 9fhs.pdb file due to purification / crystallization, the RFdiffusions were targeted to renumbered F13L chainA, to synchronize with the RFdiffusion numbering.

RFdiffusion inputs of unique peptide sizes per run (i.e., 0-, 20-, 25- and 30-mer), predicted maximal affinities for 25-mer peptides despite using different hot-spots (**Figure 1, orange squares** and **circles**). However, most of those peptides predicted only single α-helices, and many of them targeted also amino acids outside the F13L homodimer interface (not shown).

**Figure 1.**
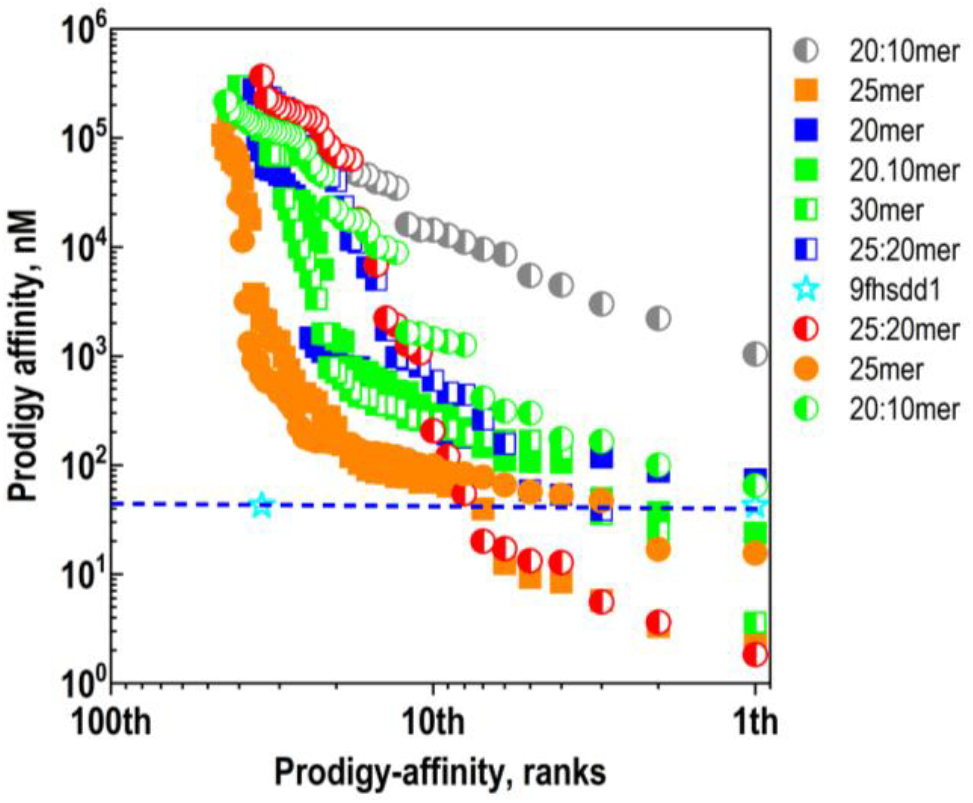
Affinity rank profiles of peptides of different sizes generated by RFdiffusion. The Prodigy affinities of 576 peptides from 9 runs complexed to the F13L chainA (9fhs.pdb) by RFdiffusion using different peptide size targets (**Table1**) were ranked. Their affinities were estimated within -- distance cutoff of 5.5 Å, using a home-designed windows-Prodigy/PyMol/Python batch-script. The top-peptide sequences predicting higher affinities (lower docking scores) than the F13L B dimer and targeting any of the two α-helices of the homodimer interface were shown **in Table 1**. See more details at **SupplementaryMaterials/Figure S1.pdb** for one run example. **Blue horizontal dashed line**, A-B chain Prodigy affinity of the 9fhs.pdb complex^14^

Some of the RFdiffusion inputs of two peptide sizes per run (i.e., combining 10-, 20- and 25-mer), generated dipeptides with high affinities (**Figure 1, half-red circles**). PyMol visual inspection of the resulting F13L chainA-dipeptide complexes predicted peptide-binder α-helices docking to the homotrimer interface (**Table 1,** 13. 7n2, 15. 12n3 and *16.9n2) and/or including terminal linear peptides extending their targeting to other amino acids (**SupplementaryMaterials/ Figure S1.pdb**).

Taking into consideration all the peptides randomly generated by several RFdiffusion runs, their corresponding ranking of Prodigy affinity profiles confirmed that most of the highest affinities were predicted when input/output contained either 25-mer peptides or 25-+ 20-mer dipeptides (**Figure 1, orange squares** and **half-red circles,** respectively).

Nine top-peptides (1.5 %) were selected for further work because: **i)** predicted =< 46 nM affinities (minimal affinity to compete with the the 9fhs.pdb AB homodimer as estimated with the same Prodigy criteria), and **ii)** interacted with α-helices of the F13L homodimer interface. The selected top-peptide-binders predicted different amino acid sequences (**Table 1, Top-peptide sequences**).

Comparison between the 9fhs.pdb homodimer chains AB model complexed with Tecovirimat and top-peptide-binders (**Figure 2**) confirmed that Tecovirimat docked to the dimer interface between α-helices (**Figure 2A, red spheres**) while most top-peptides docked to a similar site occupying the chain B interface (**Figure 2B, Different color cartoons**). These results may suggest that RFdiffusion generated top-peptide-binders could interfere with the F13L dimerization in a Tecovirimat-binding manner.

**Figure 2.**
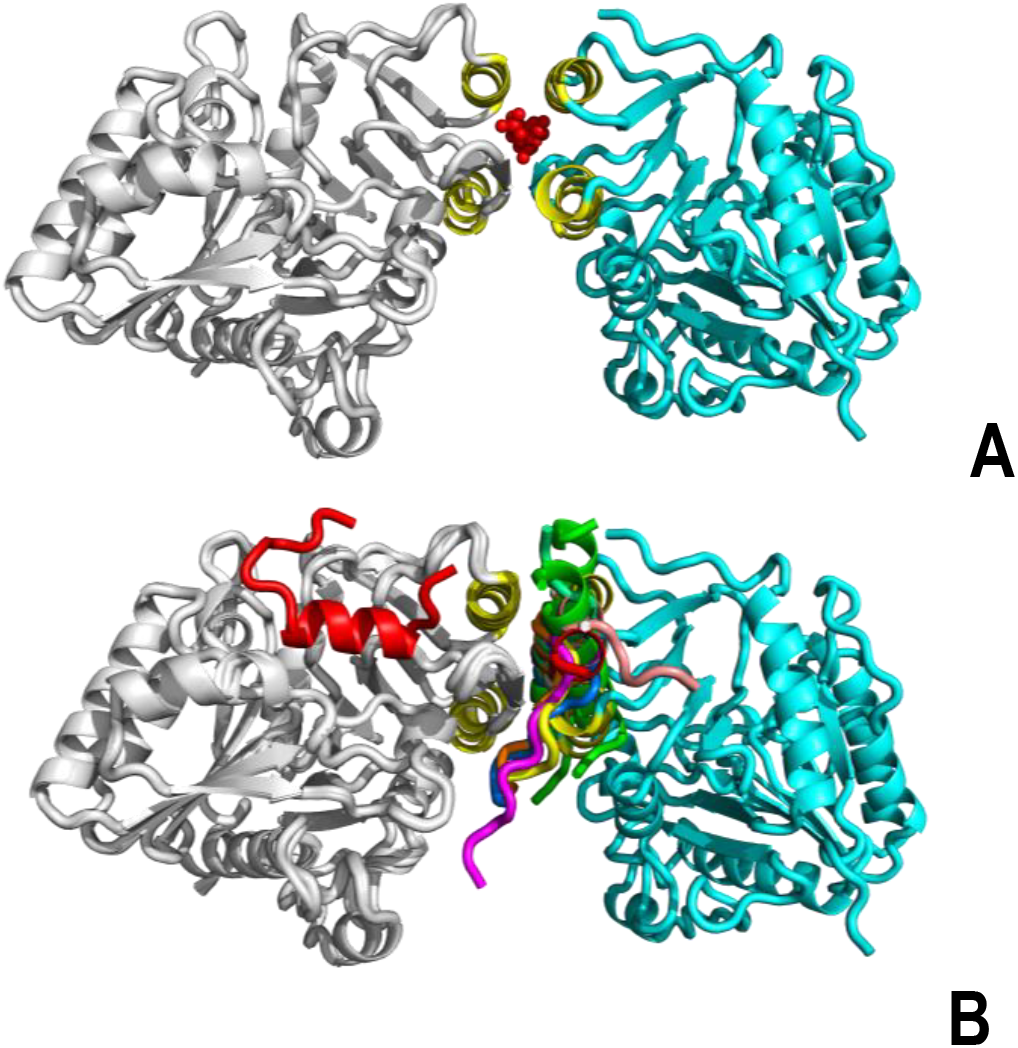
RFdiffusion models comparison between F13L homodimer and top-peptides (Table 1). The VACV F13L homodimer 9fhs.pdb was drawn as cartoons in PyMol. **A)** F13L homodimer docked to Tecovirimat. **B)** Top-peptides generated by RFdiffusion of the F13L chainA merged with the F13L chainB. **Gray,** cartoon representation of the F13L chainA **Cyan,** cartoon representation of the F13L chainB **yellow α-helices,** F13L homodimer interface α-helices (corrected numbering: 251-263+279-294) **Red spheres**, Tecovirimat docked to 9fhs.pdb by AutoDockVina in PyRx **Different color cartoons**, merged example of RFdiffusion generated top-peptides

To further clarify the top-peptide-binder targets, the F13L chainA amino acid residues nearby 5.5 Å were represented. Many of the top-peptide-binders targeted not only the amino acids included into the homodimer F13L **(Figure 3, yellow vertical rectangles**), but also its ^253^YW and phospholipase-like ^312^NxKxxxxD motifs (corrected residue positions) (**Figure 3, arrows**).

**Figure 3.**
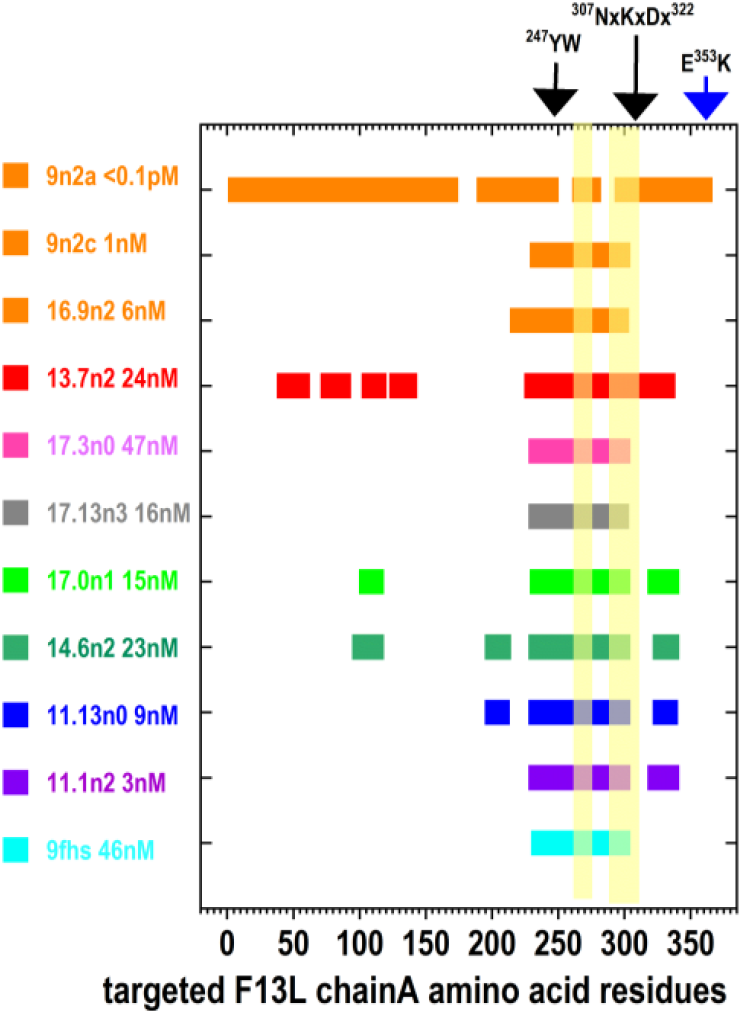
F13L chainA amino acid residues targeted by RFdiffusion top-peptides. F13L chainA amino acid residues nearby 5.5 Å to the docked F13L chainB (9fhs.pdb). The F13L chainA amino acids nearby RFdiffusion top-peptide-binders (**Table 1**) and their affinities were predicted by Prodigy. F13L mutation sensitive motif 253YW and phospholipase-like motif ^312^NxKxxxxD with corrected positions. **X- axis**, amino acid corrected positions at chainA of 9fhs.pdb (amino to carboxy-terminal residues). **Yellow vertical rectangles**, main α-helices implicated at the F13L homodimer interface (corrected positions: 251-263+279-294). The most important Tecovirimat-resistant monkeypox mutants implicated F13L residues 240-310. **9fhs.pdb**, recently solved VACV F13L dimer^14^.

Further affinity improvements in yields and affinities were obtained by including additional criteria into the RFdiffusion, such as increasing the number of random runs, fine-tuning targeted peptide sizes, number of iterations, alternative hot-spots, targeting other F13L surfaces, number of refinement cycles, genetic evolution losses algorithms, cyclization, hallucination, etc (not shown). Only some of the possible improvements by cyclization and hallucination will be shortly described now.

To improve their affinities, a first filling of gaps of dipeptide-binders were first attempted. However, in contrast to previous results ^42^, the Glycine gap-filling of the top-dipeptide-binders (**Table 1**: 13.7n2, 15.12n3 and *16.9n2), changed their targeted F13L interface elsewhere at F13L. Glycine gap-filling were not further considered for improvement optimization of the top-dipeptide-binders and therefore, alternative N-C-cyclizations were explored.

The dipeptides mentioned above re-visualized at PyMol, selected *16.9n2 as one of the higher affinities (∼ 6 nM) predicting 25- and 20-mer α-helices targeting the F13L chainA interface. Additional generations by cyclization and hallucination of random amino acid sequences were performed from the *16.9n2 peptide-binder input to generate the best top-cyclic-peptide-binders 9n2c and 9n2a (**Table 1, reddish rows**).

To generate 9n2c, the N-C cyclization was fixed to the initial *16.9n2 backbone (af_model = mk_afdesign_model (protocol=“fixbb”)). The predicted peptide sequences maintained the initial two α-helices backbone targeting the interface α-helices of F13L chainA (**Table 1, Figure 4 B, red cartoons**), increasing their predicted affinities (additional optimization results not shown). To generate 9n2a, the N-C cyclization was followed by additional sequence hallucination generations to not only different peptide sequences but also different backbones (i.e., 9n2a with a ^13^C-^26^C disulphide bond) (**Table 1, Figure 4 C, red cartoons**) targeting similar monomer inner cavities than those previously proposed by small-molecule co-evolutionary docking**^30^.**

**Figure 4.**
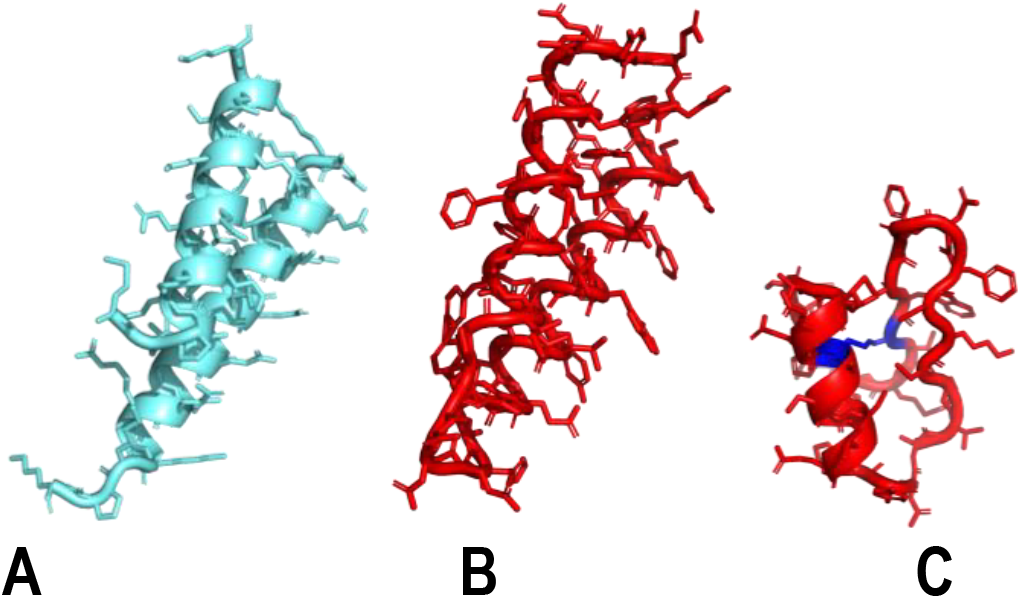
Representation of the top-peptides best-fitting the F13L dimer-interface or cavity-binding. Details of methods are provided in **Table 1** and interactions with the F13L chainA in the **GraphycalAbstract**. Because PyMol do not draws cyclization bonds on cartoons, the sticks drawings were also activated. **A cyan**, top-dipeptides 16.9n2 (25-, 20-mer), generated by RFdiffusion-main targeting the F13L dimer interface **B red**, 9n2c top-peptide designed with af_cyc_design from 16.9n2 targeting the F13L dimer interface **C red**, 9n2a top-peptide designed with af_cyc_design from 16.9n2 targeting previous F13L monomer drug-docking cavity. **C blue**, ^13^C-^26^C disulfide bond stabilizing the 9n2a structure

Since only three residues (A^72^V, C^98^S and E^353^K, original numbers) were different between F13L VACV (YP_232934) and MPXV (URK20480), comparisons of top-cyclic-peptide-binder affinities were made between VACV F13L and MPXV Tecovirimat-resistant mutants F13L0, F13L1 and F13L2, previously reported**^30^.** For that, the 9n2c and 9n2a top-cyclic-peptide-binder complexes with VACV F13L were first merged / aligned with each of the alphafold modeled MPXV F13Ls mutants**^30^**. Then, the VACV F13L sequences were deleted and Prodigy affinities the complexes between the top-cyclic-peptide-binders and MPXV F13L mutants estimated. Although preliminary, the results suggested only small affinity reductions on the mutant F13L MPXV-9n2c complexes, while the highest affinities were maintained for the F13L MPXV-9n2a complexes (**Table 2**). Ongoing results indicated that many more cyclic-peptide-binder sequence alternatives can be computationally generated predicting higher affinities by additional fine-tuning of RFdiffusion (not shown)…….

**Table 2.**
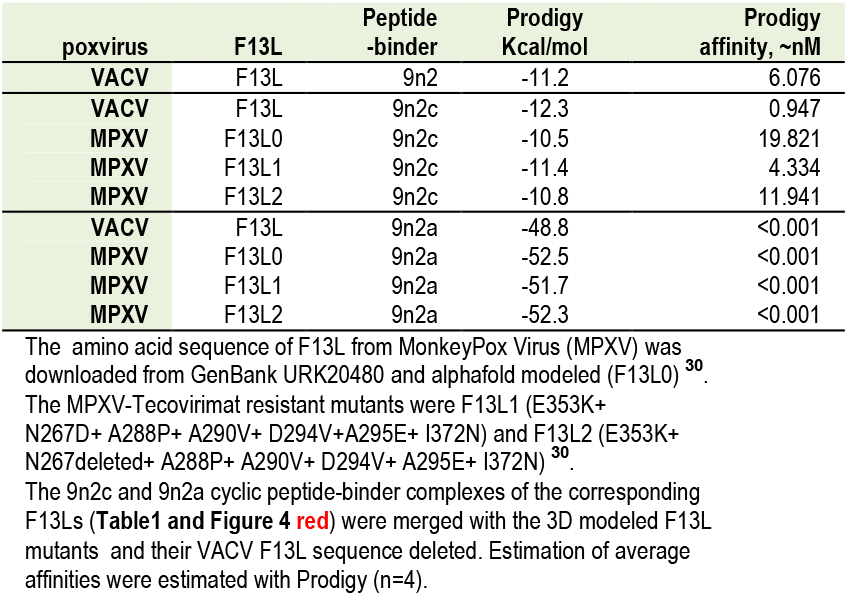
VACV and MPXV Tecovirimat resistant mutants docked to cyclic peptide-binders.

## Conclusions

Top peptide-binders predicting higher affinities to the F13L chainA interface than to chainB, have been computationally predicted by RFdiffusion-main algorithms. Additionally, top-dipeptide-binders were computationally cycled to increase their physiological stability and predicted affinity improvements and new backbones. The identification of new top-cyclic-sequences and backbones targeting F13L α-helices or previous docking cavities on VACV and MPXV for small drug-like compounds, suggest possible interferences with poxvirus homodimer formation and infection. These new cyclic-peptide-binder candidates can be further improved in their affinities but they need experimental validation after production by recombinant bacteria and/or peptide synthesis.

## Computational methods

### F13L homodimer model

The amino acid sequences corresponding to the F13L envelope phospholipase OPG057 of **Vac**cinia **V**irus (VACV) Western Reserve (695871) was downloaded with the 9fhs.pdb corresponding to the crystallizatio^n^. Because of the 32 amino acid residues containing the amino-terminal poliH tails added to the recombinant F13L to be isolated and crystallized, the targeting to most of the F13L whole amino acid sequence coded in the 9fhs.pdb required their PyMol renumbering to synchronize with the automatic renumbering the RFdiffusion algorithms will otherwise introduce. The final renumbering per F13L chain was confirmed as ^1^SVPAGA SLKI^367^. The validation of the resulting new 9fhs.pdb dimer was made by Tecovirimat docking (**A**uto**D**ock**V**ina (ADV) included into the PyRx-098/PyRx-1.0 package^46^ (https://pyrx.sourceforge.io/). As expected from the crystalized 9fhs.pdb model, Tecovirimat also docked to the interface centre of the 2×2 α-helices in the poliH-less 9fhs.pdb (**Figure, red spheres**). For RFdiffusion, the F13L chainB was removed by PyMol to yield F13L chainA.

The amino acid sequences corresponding to F13L0 from **M**onkey**P**o**x V**irus (MPXV) infection, were obtained from the 2022 international outbreak of isolate clade IIb lineage B.1 (containing the E353K mutation)^23^, was downloaded from GenBank URK20480. The F13L amino acid sequence was alphafold modeled (F13L0) (https://colab.research.google.com/github/sokrypton/ColabFold/blob/main/AlphaFold2.ipynb), using the phospholipase D of *Streptomyces sp* (ID 1v0y) as model (https://www.rcsb.org/structure/)^9^. The F13L containing the most abundant Tecovirimat-resistant mutations^23^ were Alphafold modeled as described in before^30^. The MPXV-resistant mutants contained the common *E353K* and the most abundant Tecovirimat-resistant mutations. They were **F13L1** (mutations *E353K*+N267D+ A288P+ A290V+ D294V+ A295E+ I372N) and **F13L2** (mutations *E353K*+ N267deleted+ A288P+ A290V+ D294V+ A295E+ I372N), as described before ^30^.

### *De novo* generation of peptide-binders by RFdiffusion

For **R**osetta**F**old diffusion (RFdiffusion)^43^, the renumbered VACV F13L chainA 9fhs.pdb monomer described above was targeted. RFdiffusion was trained before by noising-denoising of known protein pdb structures(Protein Data Bank) ^44 32^ (https://github.com/Rosetta-Commons/RFdiffusion).Many iterations minimizing the differences between predicted RFdiffusion structures and their initial protein structures were performed to train those noising-denoising networks. Because noising/denoising are random, different sequences can be generated from starting poly-Glycine backbone-peptides by taking into account user-provided peptide lengths, hot spots, and additional input criteria such as: number of designs=16, num_seqs=4 per design, initial-guess=OK, and num_recycles = 6. Through many iterations, the Gly positions were filled with the best-fitting amino acid residues by proteinMPNN (neural networks). Finally, to increase their prediction accuracy, the resulting peptide sequences and their 3D structures were filtered by Alphafold2 modelling^44^. The inputs required including ß-sheet enrichments and soluble peptide restrictions were:

1. The F13L chainA plus the range of their surface-contig residues (A1-367:),
2. The final number of amino acids desired for peptide-binders (either unique i.e., :30 or multiple i.e., :25:10 to explore higher affinities further).
3. one of the three lists of preferential hot-spots targeting the F13L chainA interface (short = ‘A258,A266,A285,A291,A296’; long = ‘A258,A262,A266,A284,A285,A291,A292,A296’; longer = ‘A236,A238,A239,A241,A244,A245,A246,A247,A248,A249,A250,A253,A254,A257,A258,A261,A262, A276,A277,A278,A279,A280,A283,A286,A287,A290,A291,A293,A294,A295,A296,A297,A366,A367’
4. The 9fhs.pdb target file coding for the 3D-coordinates of F13L chainA.

(https://colab.research.google.com/github/sokrypton/ColabDesign/blob/main/rf/examples/diffusion.ipynb).

During this work, 6 runs filtered 576 peptide complex outputs with averaged RMSD between chainA and peptide-binders < 50 Å. PyMol visualization selected the top-peptides from the different runs predicting interactions with the interface α-helices of chainA (**Table1**).

### Estimation of docking affinities by Prodigy

The Prodigy algorithm was used to confirm the RMSD-selected top-peptides with relative binding affinity predictions between F13L chainA and the peptide-binders^45-49^. To validate the pdb structures, calculate interatomic distances, and calculate buried surface area, Prodigy rely on Biopython https://www.biopython.org), freesasa (https://github.com/mittinatten/freesasa), and naccess (http://www.ncbi.nlm.nih.gov/pubmed/994183), respectively. Further to the Prodigy web service (https://wenmr.science.uu.nl/prodigy/), a local anaconda3-guided Prodigy program was run in Windows 10 running on Python 3.10.8. The Windows-Prodigy version was run in batch as described before^42^ through home-designed PyMol/Python for-loop scripts (subprocess.run([prodigy_exe, abs_pdb, “--selection”, “A”, “B”,

“--distance-cutoff”, “5.5”,”--contact_list”,”--pymol_selection”,”-q”], stdout=outfile, stderr= subprocess.STDOUT). The output (stdout) included all_pdb (the complex pdb files), the --distance cutoff = 5.5 Å, the contact_list (a text file of all amino acid predicting contacts), the -- pymol_selection (PyMol pml scripts drawing the side chains implicated in the contacts), and the estimated docking affinities in Kcal/mol. As recommended by the authors, to estimate the relative affinities including most weaker long protein-peptide interactions, the Prodigy affinities were calculated by a “--distance cutoff” of 5.5 Å^42^.

### Peptide cyclization

In the runs where inputs included two peptide sizes (dipeptides of), the generated peptide gaps were first filled by Glycines as detailed before^42^. The single letter amino acid code of the corresponding complexes with F13L chainA were submitted to Alphafold-batch (https://colab.research.google.com/github/sokrypton/ColabFold/blob/main/batch/AlphaFold2_batch.ipynb), through Google Drive/input_batch directory containing the fasta files. In the batch notebook version, the results were automatically delivered to the Drive, locally downloaded and saved to an all_pdb directory by a home-designed PyMol/Python script. All the modeled *.pdbs were finally submitted to Prodigy as mentioned above to analyse the results. In this work the filling gaps with polyGly changed the targeted F13L site and therefore were considered not improved and to conserve their targeting cyclization was attempted.

The predicted complexes with dipeptides visualized at PyMol, selected the 16.9n2 complex as one of the higher affinities (∼ 6 nM) plus predicting 2 peptide α-helices of 25- and 20-mer which targeted amino acids nearby the two interface α-helices. Cyclization and hallucination (generation) of additional amino acid sequences were performed in the 16.9n2 peptide-binders to generate the best α-helix peptide sequences 9n2c and 9n2a (**orange rows**) using the Colab notebook: https. The 9n2c cyclic peptide-binder was the best predicted peptide sequence with one backbone and target very similar to the initial 9n2 dipeptide-binder sequence obtained by using the fixed backbone design (af_model = mk_afdesign_model(protocol=“fixbb”)).

**The 9n2a peptide cyclic peptide-binder,** was the best predicted peptide sequence with a different backbone targeting the previously inner monomer cavity identified by docking obtained by including not only cyclization but also amino acid hallucinations (generations) of the initial 9n2 dipeptide sequence. One disulphide bond was predicted between the ^13^C and ^26^C of 9n2a (**red carbons**).

## Supplementary information

**Table S1.**
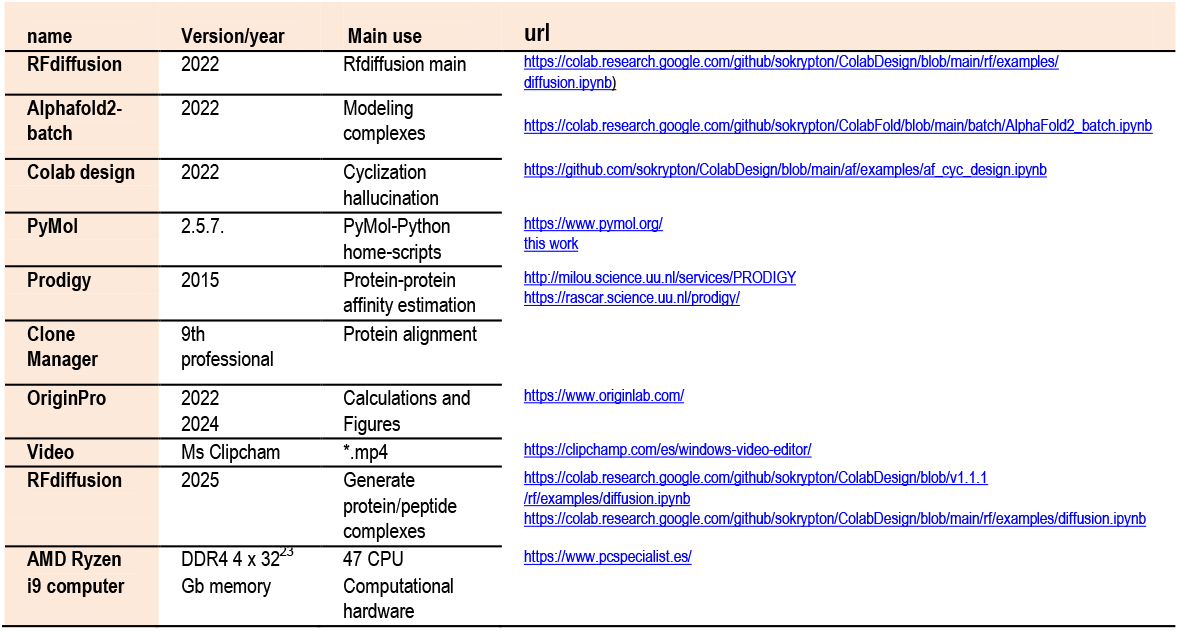
Computational software and hardware.

### Supplementary Materials

**GraphycalAbstract.pdb. Grey cartoon**, carbon backbone of VACV F13L chainA (9fhs.pdb) ^14^. **yellow cartoons,** two α-helix sequences most implicated in the F13L homodimer interface (renumbered to 251-263+279-294). **Ligh green α-helices**, 25- and 20-mer dipeptide-binder 16.9n2 α-helixes generated by RFdiffusion and used as fixed backbone (fixbb) for cyclization (**Table1**). **Red cartoon right,** fixbb cyclization maintaining backbone and interface targeting (9n2c). **Red cartoon left,** cyclization + hallucination generating different sequence / backbone and targeting the inner F13L previously identified by co-evolutionary docking**^30^**(9n2a). **Red sticks** were added to the red cartoons to show their N-C cyclization bonds at PyMol. **Blue spheres,** 9n2a ^13^C - ^26^C internal disulphide bond.

**Figure S1.pdb. Example of RFdiffusion designs. Grey cartoon,** carbon backbone of VACV F13L chainA (9fhs.pdb) ^14^. **yellow cartoons**, two α-helix sequences most implicated in the F13L homodimer interface (renumbered to 251-263+279-294). All_pdb results of one RFdiffusion run targeting VACV F13L chainA 1-367, 25- and 20-mer contigs, to randomly generate 16 designs, 4 sequences per design, including ß-sheet and soluble peptide preferences. **Different colour cartoons,** RFdiffusion-generated peptides.

## Supporting information

Graphycal Abstract

## Competing interests

The author declares no competing interests

### Authors’ contributions

JC designed, performed and analyzed the computational work and drafted the manuscript.

## Acknowledgements

Thanks are due to Dr. Bonwin M.J.J. of Bijvoet Center for Biomolecular Research, Utrech, The Netherlands for his initial help with the Prodigy installation at Windows.

